# Notch signalling influences cell fate decisions and HOX gene induction in axial progenitors

**DOI:** 10.1101/2023.06.16.545269

**Authors:** Fay Cooper, Celine Souilhol, Scott Haston, Shona Gray, Katy Boswell, Antigoni Gogolou, Thomas Frith, Dylan Stavish, Bethany M James, Dan Bose, Jacqueline Kim Dale, Anestis Tsakiridis

## Abstract

The generation of the post-cranial embryonic body relies on the coordinated production of spinal cord neurectoderm and presomitic mesoderm cells from neuromesodermal progenitors (NMPs). This process is orchestrated by pro-neural and pro-mesodermal transcription factors that are co-expressed in NMPs together with Hox genes, which are critical for axial allocation of NMP derivatives. NMPs reside in a posterior growth region, which is marked by the expression of Wnt, FGF and Notch signalling components. While the importance of Wnt and FGF in influencing the induction and differentiation of NMPs is well established, the precise role of Notch remains unclear. Here, we show that the Wnt/FGF-driven induction of NMPs from human embryonic stem cells (hESCs) relies on Notch signalling. Using hESC-derived NMPs and chick embryo grafting, we demonstrate that Notch directs a pro-mesodermal character at the expense of neural fate. We show that Notch also contributes to activation of *HOX* gene expression in human NMPs, partly in a non cell-autonomous manner. Finally, we provide evidence that Notch exerts its effects via the establishment of a negative feedback loop with FGF signalling.

## INTRODUCTION

The formation of the amniote embryonic body takes place in a head-to-tail (anterior-posterior) direction and it is driven by developmentally plastic axial progenitors, which can generate both spinal cord neurectoderm and presomitic/paraxial mesoderm, the precursor of the vertebral column/trunk musculature (thus termed NMPs; reviewed in (Wymeersch et al., 2021)). NMPs arise around the end of gastrulation/early somitogenesis, within a posterior growth region that encompasses the node-anterior primitive streak border (NSB) and the caudal lateral epiblast (Brown & Storey, 2000; Cambray & Wilson, 2002, 2007; Guillot et al., 2021; Mugele et al., 2018; Wymeersch et al., 2016). They are marked by the co-expression of pro-neural and pro-mesodermal transcription factors, such as *Sox2, T/Brachyury* (*TBXT* in humans), *Tbx6* and *Cdx2* (Gouti et al., 2017; Guillot et al., 2021; Javali et al., 2017; Koch et al., 2017; Martin & Kimelman, 2012; Olivera-Martinez et al., 2012; Tsakiridis et al., 2014; Wymeersch et al., 2016). The antagonistic interaction between these lineage-specific transcription factors determines the balanced production of neural vs mesodermal cell types from NMPs (Gouti et al., 2017; Koch et al., 2017). NMPs are also marked by the expression of *Hox* gene family members (arranged as paralogous groups [PG] in four distinct chromosomal clusters: A, B, C, and D), which are activated within the posterior growth region in a sequential manner reflecting their 3’-to-5’ genomic order (Gouti et al., 2017; Guillot et al., 2021; Neijts et al., 2017; Wymeersch et al., 2019). The latter process is tightly linked to the assignment of a positional identity in the nascent axial progenitor derivatives before their allocation along the developing embryonic anteroposterior axis (reviewed by (Deschamps & Duboule, 2017)).

The NMP niche relies on the activity of key posteriorizing signalling pathways, such as Wnt and FGF. These trigger the transcription factor networks operating within NMPs, which in turn, potentiate, via positive feedback, Wnt/FGF activity within the posterior growth region during axis elongation (Amin et al., 2016; Blassberg et al., 2022; Martin & Kimelman, 2012; Mukherjee et al., 2022; Young et al., 2009). The balance between these two signalling pathways appears to orchestrate NMP cell fate decisions as Wnt/FGF have been shown to be linked to both progenitor maintenance and differentiation toward early neural and presomitic mesoderm cells (Amin et al., 2016; Anand et al., 2023; Cooper et al., 2022; Delfino-Machín et al., 2005; Diez del Corral et al., 2002; Gouti et al., 2017; Martin & Kimelman, 2012; Semprich et al., 2022; Wind et al., 2021; Young et al., 2009). In line with these findings, Wnt and FGF signalling agonists are the two main components of protocols for the generation of NMP-like cells and their earliest mesodermal and neural derivatives from mouse and human pluripotent stem cells *in vitro* (Chal et al., 2015; Cooper et al., 2022; Frith et al., 2018; Lippmann et al., 2015; Turner et al., 2014; Verrier et al., 2018; Wind et al., 2021). Moreover, *Hox* gene expression in the posterior growth region/NMPs is also driven largely by Wnt and FGF activity via crosstalk with the two key posteriorizing transcription factors *CDX2* and *TBXT* (Amin et al., 2016; Chawengsaksophak et al., 2004; Gogolou et al., 2022; Metzis et al., 2018; Neijts et al., 2017; Neijts et al., 2016).

The other key developmental signalling pathway that has been found to be active in the posterior growth region/NMP niches is Notch. Notch signalling is activated through the interaction of receptors and ligands expressed by neighbouring cells. In mammals, there are four transmembrane receptors (NOTCH 1-4), which bind to five NOTCH transmembrane ligands (DLL1, DLL3, DLL4, JAG1 and JAG2). Once bound, the NOTCH receptor undergoes two successive proteolytic cleavage events mediated by ADAM10 and γ-SECRETASE which releases the intracellular NOTCH domain (NICD) into the cell nucleus and allowing it to bind to the NOTCH signalling transcription factor RBPJk/CSL (Carrieri & Dale, 2016; Shen et al., 2021). Several Notch signalling components are expressed in NMPs and their immediate neural and mesodermal derivatives, from late gastrulation and throughout embryonic axis elongation (Akai et al., 2005; Bettenhausen et al., 1995; Dunwoodie et al., 1997; Williams et al., 1995; Wymeersch et al., 2019; Zhang & Gridley, 1998). Moreover, the attenuation or overexpression of many of these components leads to severe posterior patterning defects (Akai et al., 2005; Dale et al., 2003; de la Pompa et al., 1997; Donoviel et al., 1999; Nowotschin et al., 2012; Oka et al., 1995; Souilhol et al., 2015). Notch signalling has also been found to crosstalk with the principal posteriorizing Wnt and FGF signalling pathways during axis elongation (Akai et al., 2005; Galceran et al., 2004; Gibb et al., 2009; Nakaya et al., 2005). and the expression of Notch signalling components in the posterior growth region is driven by key NMP regulators-Wnt/FGF targets such as *T/TBXT* and *Cdx2* (Amin et al., 2016; Gogolou et al., 2022; Guibentif et al., 2021; Koch et al., 2017). Collectively, these data suggest that Notch signalling may be a critical component of the NMP niche and interlinked with the well-established signalling pathways regulating NMP specification and maintenance. However, it is still unclear how exactly Notch influences NMP ontogeny.

Here, we investigated the role of Notch signalling in axial progenitors using the differentiation of human embryonic stem cells (hESCs) toward NMPs as a model. We show that Notch attenuation during NMP induction impairs the activation of pro-mesodermal transcription factors and global *HOX* activation whilst promoting an early neural character. Our results indicate that Notch-driven pro-mesodermal/*HOX* gene expression control is mediated via the establishment of a feedback loop with FGF signalling. We provide evidence that the induction of certain *HOX* genes in hESC-derived NMPs may be mediated by Notch in a non-cell autonomous fashion. Finally, Notch signalling inhibition in chick embryonic NMPs dramatically alters their engraftment behaviour and impairs their capacity to generate paraxial mesoderm cells biasing them instead toward a ventral neural/floor plate cell fate. Together, these findings suggest that Notch contributes, together with Wnt and FGF, to the primary signalling axis within the posterior growth region that orchestrates NMP cell fate decisions and positional identity acquisition.

## RESULTS AND DISCUSSION

### Notch signalling mediates the induction of pro-mesodermal and *HOX* genes in NMPs

We have previously shown that the *in vitro* generation of NMPs following treatment of hPSCs with the Wnt agonist CHIR99021 (CHIR) and recombinant FGF2 for three days is accompanied by an upregulation of Notch signalling-associated transcripts (Frith et al., 2018; Wind et al., 2021), in line with findings demonstrating high Notch activity in the early posterior growth region and NMPs around the end of gastrulation/early somitogenesis *in vivo* (Bettenhausen et al., 1995; Dunwoodie et al., 1997; Williams et al., 1995; Wymeersch et al., 2019). To interrogate the role of this increase in Notch signalling activity during the transition of pluripotent cells toward a neuromesodermal-potent state, we generated NMPs from WA09 (H9) hESCs in the presence of the Notch/γ-secretase inhibitor DAPT or DMSO (control) (**Fig. 1A**). Quantitative PCR (qPCR)-based analysis of DAPT-treated NMP cultures (NOTCHi) revealed that they expressed significantly reduced levels of Notch target genes/components, particularly *HES5,* compared to controls, indicating effective attenuation of Notch signalling (**Fig. S1A**). Moreover, NOTCHi NMPs were marked by a considerable reduction in the expression of pro-mesodermal/NMP markers such as *TBXT*, *TBX6* and *CDX1* and a concomitant increase in the transcription of the pro-neural NMP marker *SOX2* (**Fig. 1B**). Similar changes in TBXT and SOX2 were detected at the protein level (**Fig. 1C, D**), while we found no increase in the expression of pluripotency-associated (OCT4 and NANOG) or later spinal cord neurectodermal (PAX6 and SOX1) markers, which remained low/undetected (**Fig. S1B** and data not shown). Together, these results suggest that NOTCH signalling mediates the pro-mesodermal character of NMPs during their specification from pluripotent cells at the expense of a spinal cord pre-neural SOX2+ identity.

**Figure 1.**
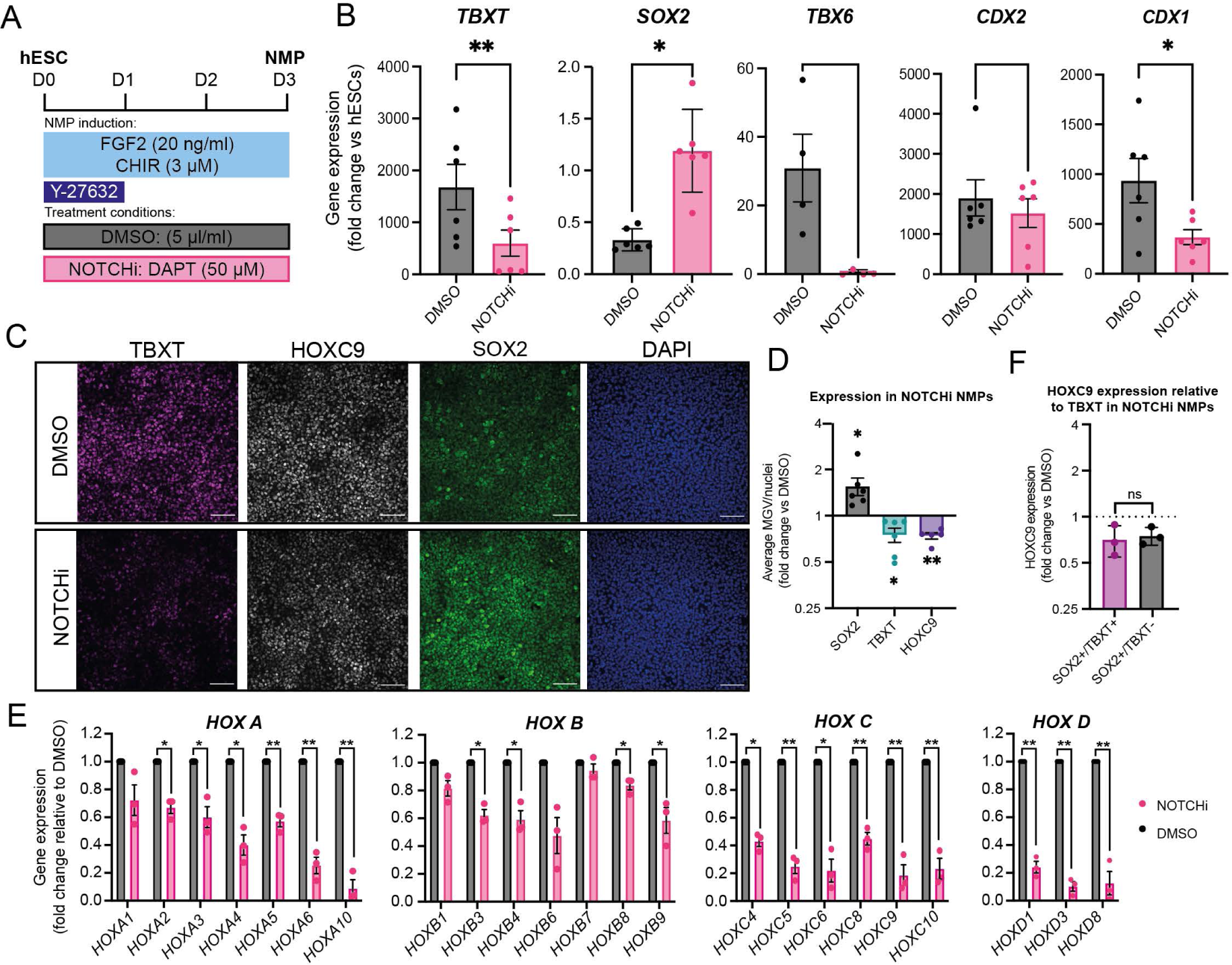
Notch inhibition impairs the induction of pro-mesodermal/*HOX* genes during NMP specification *in vitro*. (A) Schematic representation of the treatment conditions used to generate NOTCHi or DMSO control NMPs from hESCs. (B) qPCR expression analysis of key NMP markers in hESC-derived NOTCHi/control NMPs. Error bars represent s.e.m. (n=3-4). *SOX2*\*P=0.01; *TBXT*\*\*P=0.0024 and CDX1*P=0.0148 (paired t-test). (C) Immunofluorescence analysis of the expression of HOXC9, TBXT and SOX2 in NMPs treated with DMSO or DAPT. Scale bars = 100μm. (D) Image analysis of average mean gray value (MGV) per nuclei (displayed as fold change over DMSO control) of TBXT, SOX2 and HOXC9 protein expression in DAPT treated NMPs. Error bars represent s.e.m. (n=5-6). SOX2 *P=0.0413, HOXC9 **P=0.0017, TBXT *P=0.0263 (one sample t and Wilcoxon test). (E) qPCR expression analysis of indicated *HOX* genes in hESC-derived NOTCHi/control NMPs. Error bars represent s.e.m. (n=3). *P≤0.05, **P≤0.01 ***P=≤0.001 (one sample t and Wilcoxon test). (F) Immunofluorescence analysis of MGV of HOXC9 and SOX2 protein expression relative to TBXT positivity (TBXT+ or TBXT-) in DAPT-treated NMPs. Error bars represent s.e.m. (n=3) (paired t-test). ns, not significant.

We next examined the global activation of *HOX* genes, a major hallmark of Wnt/FGF-driven acquisition of a posterior axial and NMP identity (Cooper et al., 2022; Gogolou et al., 2022; Gouti et al., 2017; Guillot et al., 2021; Wymeersch et al., 2019), in DAPT-treated cultures. We found that NOTCHi hESC-derived NMPs exhibited a marked reduction in the expression of most *HOX* PG members examined, particularly those belonging to the *HOXC* and *HOXD* clusters, compared to the DMSO controls (**Fig. 1E**). Similarly, immunofluorescence analysis of NOTCHi NMP cultures revealed a decrease in HOXC9, TBXT and SOX2 protein levels relative to their DMSO-treated counterparts (**Fig. 1C, D**). This DAPT-driven perturbation in HOXC9 expression was detected in SOX2-positive/TBXT-positive as well as SOX2-positive/TBXT-negative cell populations (**Fig. 1F**) suggesting that impaired activation of *HOX* gene clusters occurs irrespectively of the expression status of TBXT, a transcription factor that has been found to control directly *HOX* gene transcription in human NMPs (Gogolou et al., 2022). Together, these findings indicate that, Notch signalling modulates the induction of a posterior axial identity and colinear activation of *HOX* PG family members by Wnt and FGF, as pluripotent cells transit toward NMPs.

### Non-cell autonomous control of *HOX* gene expression in human NMPs is partly Notch-driven

The striking effect of DAPT on the induction of various *HOX* genes in hESC-derived NMPs prompted us to further examine the links between Notch and *HOX* expression control. Heterochronic grafting experiments have indicated that the global *Hox* gene expression profile of axial progenitors is plastic as it can be ‘reset’ in response to extrinsic cues emanating from the NMP niche (McGrew et al., 2008). We have also previously shown that hESC-derived NMPs, in which *TBXT* is knocked down via a Tetracycline (Tet)-inducible, short hairpin RNA (shRNA)-mediated system (Bertero et al., 2016) (TiKD) are marked by reduced Notch activity as well as an inability to induce properly *HOX* PG(1-9) members (Gogolou et al., 2022). Given that Notch signalling is typically encoded via receptor-ligand interaction between neighbouring cells, we tested whether it could influence/rescue *HOX* gene expression in a non-cell autonomous manner. To this end, we mixed TiKD hESCs with isogenic wild type hESCs constitutively expressing an red fluorescent protein reporter (H9-RFP), at a 50:50 ratio. The co-cultures were differentiated toward NMPs and treated with Tet to mediate *TBXT* knockdown specifically in the unlabelled TiKD fraction, in the presence or absence of DAPT (**Fig. 2A**). Following NMP differentiation, TBXT knockdown/RFP-negative cells were FACS-sorted from the co-cultures and the levels of *HOX* transcripts were assayed by qPCR and compared to +/- Tet NMPs derived from TiKD hESCs without co-culture (**Fig. 2A, S2**). We found that Tet-induced *TBXT* knockdown was efficient in TiKD cells cultured either alone or together with their wild type counterparts (**Fig. 2B**). Tet-induced TBXT knockdown triggered a significant decrease in the expression of most *HOX* genes and the Notch target *HES5* (**Fig. 2C-F**, compare black vs light blue bars) as previously reported (Gogolou et al., 2022). Strikingly, this trend was partially reversed in TiKD cells upon co-culture with H9-RFP cells: the expression of some *HOX* genes, particularly those belonging to the *HOXB* PG (5-9), was restored back to levels similar to the -Tet controls (**Fig. 2C-F**, compare black vs light blue vs purple bars). Moreover, upon co-culture with H9-RFPs, TiKD cells exhibited a large increase in the levels of *HES5* (above the -Tet control levels) suggesting that Notch overactivation takes place specifically under these conditions (**Fig. 2B**, compare black vs light blue vs purple bars). As expected, this was counteracted by DAPT treatment (**Fig. 2B**, compare purple vs pink bars), which simultaneously appeared to prevent, mainly in *HOXB* cluster members, the gene expression compensatory effect of the co-culture on TiKD NMPs (**Fig. 2C-F**, compare purple vs pink bars). Co-culture/DAPT treatment did not alter the expression of TBXT relative to the Tet-treated TiKD cells cultured alone (**Fig. 2B**, compare black vs light blue vs purple vs pink bars). Collectively, these results suggest that Notch signalling can control the expression of at least a fraction of the *HOX* genes expressed by NMPs in a non-cell autonomous manner and TBXT-independent manner.

**Figure 2.**
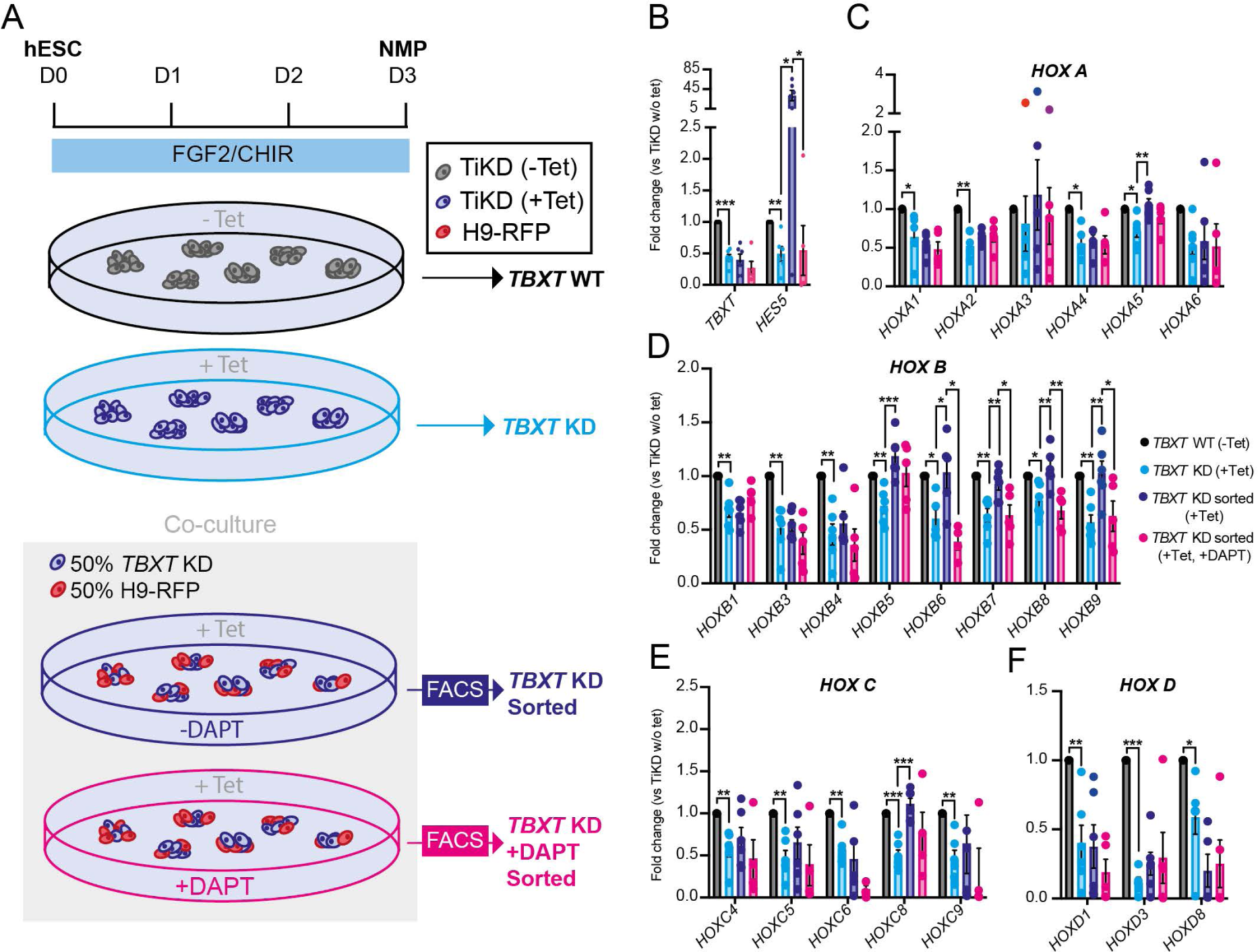
Notch signalling-dependent rescue of *HOX* gene expresion in TBXT-depleted NMPs. (A) Scheme depicting the experimental design of the *TBXT* shRNA-wild type NMP co-culture experiment. (B-F) qPCR expression analysis of *TBXT* and *HES5* (B) and *HOX* genes belonging to different paralogous groups (C-F) under the different experimental conditions depicted in A. Error bars represent s.e.m (n=3-6) *P≤0.05, **P≤0.01 ***P=≤0.001 (one sample t and Wilcoxon test (TiKD w/o Tet vs TiKD (+Tet)) or an unpaired t.test (TiKD (+Tet) vs TiKD sorted (+tet) vs TiKD (+DAPT +Tet)).

### Notch amplifies FGF activity in NMPs

To further understand how Notch signalling influences NMP specification/*HOX* gene expression, we assessed its crosstalk with the two key posteriorising signalling pathways driving embryonic axis elongation, Wnt and FGF. Thus, we generated NMPs from hESCs in the presence of either DAPT or DMSO as described above (**Fig. 1A, 3A**) and assessed the expression of Wnt/FGF signalling pathway components by qPCR. The transcript levels of Wnt target genes such as *AXIN2*, *LEF1* and *TCF1* remained unchanged in NOTCHi conditions, whereas expression of *SPRY4*, a FGF signalling target gene, was diminished (**Fig. 3B**), indicating that Notch inhibition results in a reduction of FGF signalling activity. To further confirm this, we examined the levels of the phosphorylated FGF effector kinase ERK1/2 (MAPK) by Western blot (**Fig. 3C**). Both phosphorylated p44 and p42 versions were reduced in NOTCHi NMPs compared to the DMSO-treated controls (**Fig. 3C, D**) further supporting the notion that Notch positively regulates FGF signalling in hESC derived NMPs. We further tested this, by examining whether the NOTCHi NMP phenotype can be rescued by boosting FGF signalling levels via an increase in FGF2 levels. We found that doubling the dosage of FGF2 from 20 to 40 ng/ml, in the presence of DAPT, during NMP induction from hESCs, blunted the impact of Notch inhibition and led to an increase in the expression of *TBXT* and all *HOX* genes examined back to levels comparable to those in the DMSO controls (**Fig. 3E**). Conversely, differentiation of hESCs toward NMPs in the absence of FGF2 and presence of the FGF pathway-MEK1/2 inhibitor PD0325901 (PD03) and CHIR alone (FGFi) appeared to phenocopy the effects of NOTCHi: qPCR analysis of the resulting cultures revealed the downregulation of pro-mesodermal (e.g. *TBX6*) and upregulation of pro-neural transcript (*SOX2*) (**Fig. 3F**). Unlike NOTCHi, definitive neuroectoderm genes PAX6 and SOX1 were found to be significantly upregulated in FGFi conditions (**Fig. 3F**). The expression of the FGF targets *SPRY2* and *SPRY4,* was robustly reduced confirming efficient FGF signalling inhibition under these conditions (**Fig. 3G**). FGF inhibition also resulted in a reduction of Wnt signalling components in line with findings from analysis of the embryonic NMP niches (Oginuma et al., 2017; Olivera-Martinez et al., 2012). Collectively, our data, combined with our previous observations showing that CHIR-PD03-treated hESC-derived NMPs are marked by global reduction of *HOX* gene expression as well as *TBXT* (Gogolou et al., 2022), strongly suggest that Notch signalling promotes the induction of these genes via its, direct or indirect, crosstalk with FGF signalling. Interestingly, FGF inhibition also led to a dramatic increase in the levels of the Notch target *HES5* (**Fig. 3G**), consistent with a possible feedback loop between Notch and FGF signalling (**Fig. 3H**).

**Figure 3.**
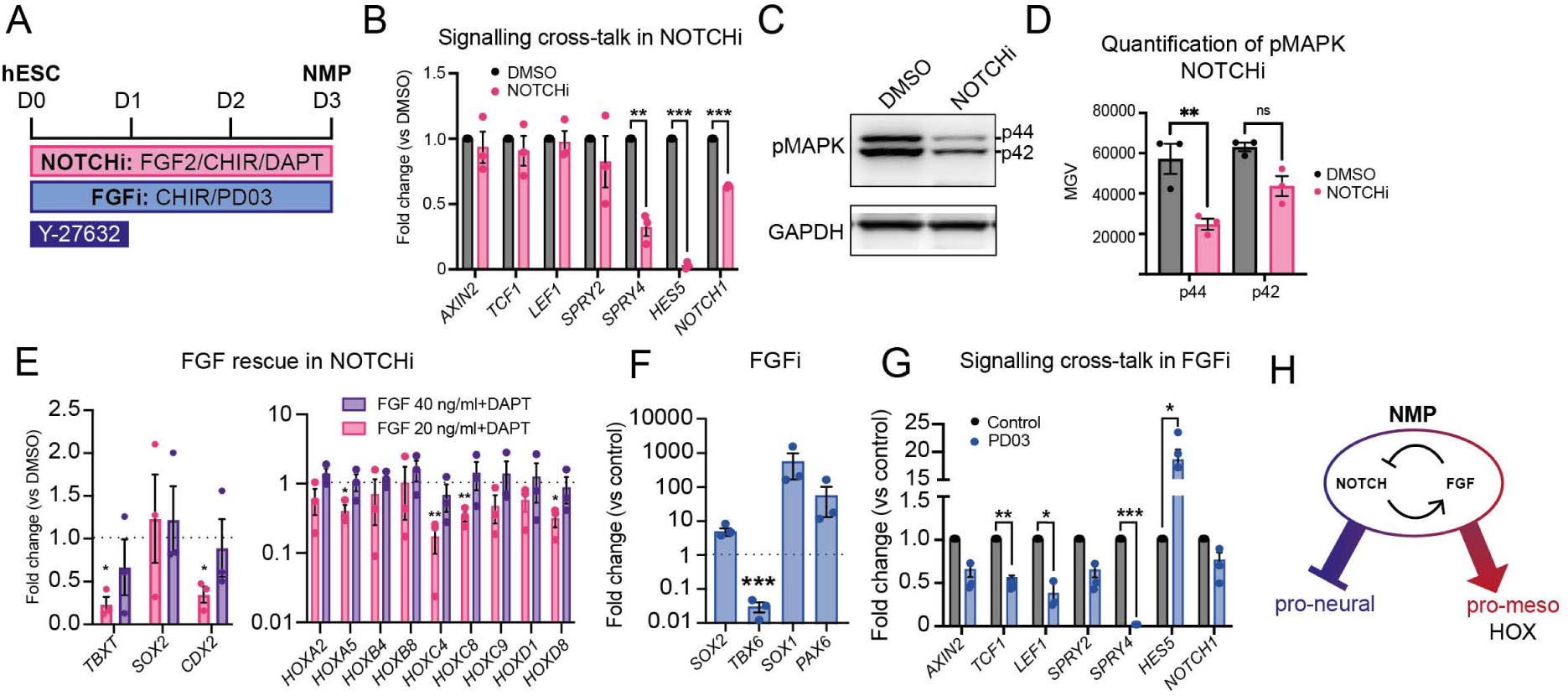
Notch-FGF signalling crosstalk in hESC-derived NMPs. (A) Scheme of treatments during the differentiation of hESCs toward NMPs. (B) qPCR expression analysis of indicated Wnt, FGF and Notch signalling pathway components in DAPT/DMSO-treated hESC-derived NMP cultures. Error bars represent s.e.m. (n=3) *SPRY4* **P=0.009, HES5 ***P=0.0002, *NOTCH1* ***P=0.001 (one sample t and Wilcoxon test). (C) Representative western blot analysis of phospho-MAPK (p42/p44) in NOTCHi/DMSO-treated NMPs and corresponding quantification (D). Error bars represent s.e.m. (n=3) p42 *P=0.036 and p44 *P=0.031 (paired t-test). (E) qPCR expression analysis of NMP markers and indicated *HOX* genes in NOTCHi NMPs generated using the standard (20ng/ml) or high (40ng/ml) FGF2 concentration from hESCs. Error bars represent s.e.m. (n=3) *P≤0.05, **P≤0.01 ***P=≤0.001 (one sample t and Wilcoxon test). (F) qPCR expression analysis of indicated pro-neural/mesodermal NMP and spinal cord neurectoderm markers in PD03-treated (FGFi) hESC-derived NMPs vs controls. Error bars represent s.e.m. (n=3) *TBX6* ***P=0.001 (one sample t and Wilcoxon test). (G) qPCR expression analysis of indicated Wnt, FGF and Notch signalling pathway components in PD03-treated/control hESC-derived NMP cultures. Error bars represent s.e.m. (n=3). *TCF* **P=0.0027, *LEF1* *P=0.0346, *SPRY4* ***P<0.0001, HES5 *P=0.01 (one sample t and Wilcoxon test).

### Notch controls axial progenitor cell fate decisions *in vivo*

We next examined the role of Notch signalling in NMP differentiation *in vivo*. To this end, wildtype and transgenic chicken embryos ubiquitously expressing green fluorescent protein (GFP) were incubated until Hamburger Hamilton (HH) (Hamburger & Hamilton, 1951) stage 4 and then dissected from the egg and cultured *in vitro* until HH8, i.e. the time window that coincides with the emergence of NMPs in the posterior growth region (Guillot et al., 2021)(**Fig. 4A**). Embryos were cultured on media plates containing either the γ-secretase Notch inhibitor LY411575 (LY) (Wong et al., 2004) or DMSO (control). Following *in vitro* culture, the NSB region from DMSO or LY-treated HH8 GFP transgenic donor chicks was isolated and grafted to a homotopic location on stage matched, DMSO or LY-treated wild type host embryos respectively (**Fig. 4A**). The host embryos were returned to their respective *in vitro* culture plates (LY or DMSO) and allowed to develop for a further 27 to 29 hours to allow for progenitor cells within the NSB to contribute to axial and paraxial tissues (**Fig. 4B)**. The contribution of GFP+ donor cells along the axis was then scored according to their final anteroposterior location and subdivided into four domains: rostral, middle, caudal and pre-progenitor (see a-e in **Fig. 4B**).

**Figure 4.**
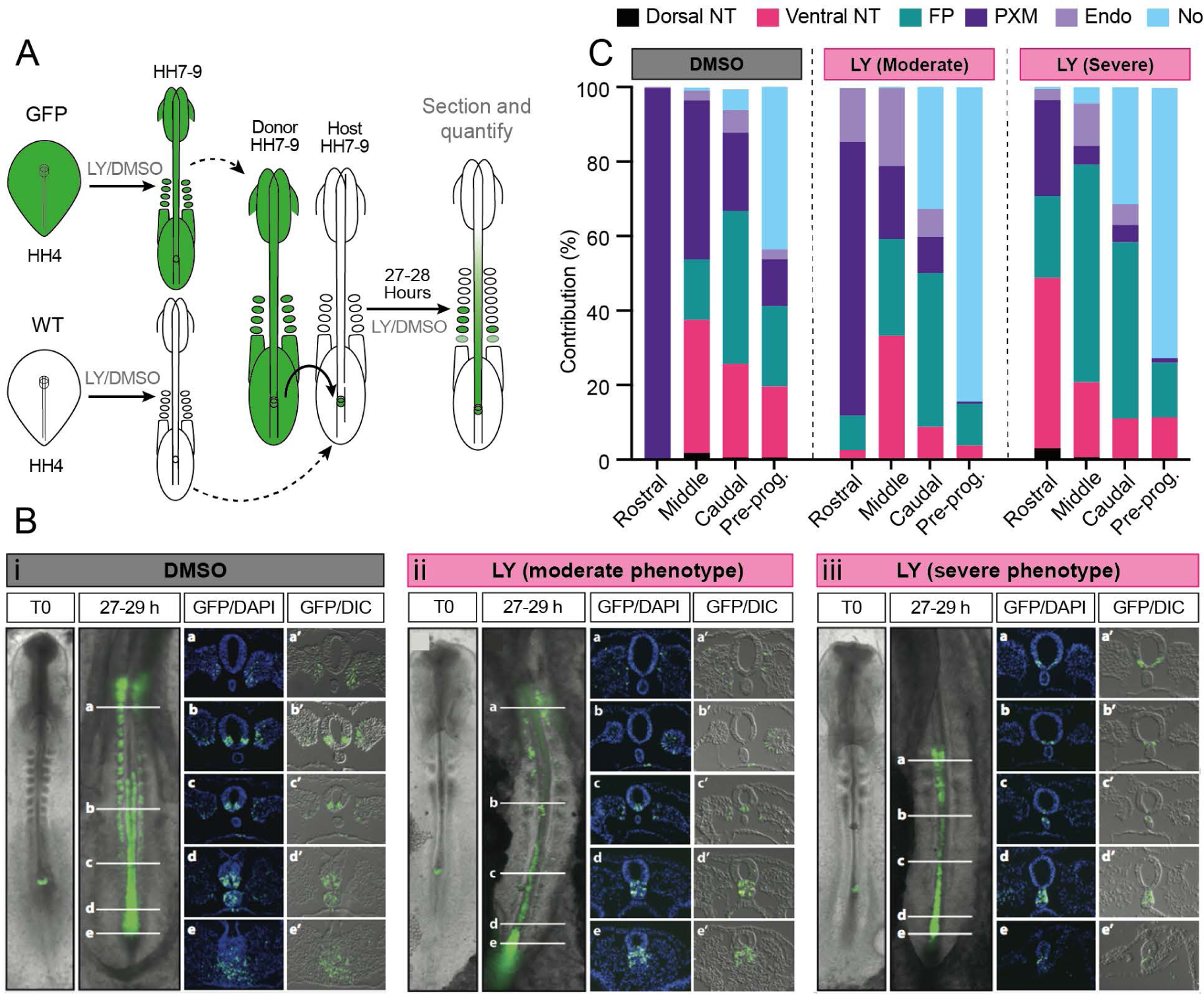
Notch signalling influences the contribution profile of axial progenitor cells *in vivo*. (A) Scheme depicting the experimental design/treatment regimens of chick embryo grafting experiments. (B) Wholemount embryo at the time of receiving a NSB graft (T0) and the GFP contribution pattern following culture in the presence of the (i) DMSO or the Notch inhibitor LY in both the moderate (ii) and severe (iii) embryos after 27-29 hours following the graft. Transverse sections at the level of the white indicator lines (a, b, c, d, e) show the nuclear stain DAPI and GFP or DIC with GFP (a’, b’, c’, d’, e’). Images are representative of independent experiments (analysed sectioned embryos: DMSO n=9, LY severe n=4/9 and moderate n=5/9). (C) Quantification of the proportion (%) of GFP cells contributing to axial and paraxial structures (dorsal neural tube (dorsal NT), ventral neural tube (ventral NT), floor plate (FP), paraxial mesoderm (somites rostrally and PSM caudally, PXM), endoderm (Endo) and the notochord (No)) in (i) DMSO and (ii) LY-treated embryos.

Fluorescence microscopy analysis of grafted host embryos revealed that in both DMSO (n=9) and LY treatment (n=13) conditions the extent of donor cell contribution along the anteroposterior axis was similar (**Fig. S3A**). We found that in the case of DMSO-treated embryos, GFP labelled donor axial progenitors contributed almost exclusively to paraxial mesoderm (PXM, >99%) in the rostral domain whereas in the more posterior domains (middle, caudal and pre-progenitor), GFP+ cells were detected in both PXM and the ventral/floor plate segments of the neural tube (ventral NT and FP respectively; **Fig. 4B, C, S3B**) denoting the NM bipotency of the grafted donor NSB fragments. The contribution of the donor cells to the dorsal neural tube in the middle, caudal and pre-progenitor domains was minimal while the number of donor cells in the notochord (No) increased in an anterior-posterior direction (**Fig. 4B, C, S3B**; n=9). These findings are in line with previous studies demonstrating the presence of ventral NT/FP/notochord-biased axial progenitors located in the early somite-stage NSB/node in amniote embryos (Cambray & Wilson, 2007; Catala et al., 1996; Mugele et al., 2018; Selleck & Stern, 1991; Wilson & Beddington, 1996; Wymeersch et al., 2016). We also detected a few GFP+ cells in the gut within the caudal/pre-progenitor (anterior streak) domains, likely reflecting the inclusion of early node or anterior primitive streak-located endoderm progenitors (“Endo”, **Fig. 4B, C, S3B**) (Selleck & Stern, 1991; Wilson & Beddington, 1996). In contrast, the most severely affected LY-treated embryos (“severe”; n=4/9) exhibited very little/no PXM contribution of GFP+ donor cells, whose presence was mainly confined to the FP and to a lesser extent the ventral NT as well as the notochord in the caudal/pre-progenitor domains **(Fig. 4B, C, S3B**). A second class of LY-associated “moderate” (n=5/9) phenotype embryos displaying intermediate features between the DMSO and severe LY treatments was also identified **(Fig. 4B, C, S3B**). Collectively, these findings suggest that Notch signalling preferentially biases NSB-located NMPs to contribute to the paraxial mesodermal lineage at the expense of a ventral neural tube/floor plate fate.

In summary, here we demonstrate that Notch is a central component of the signalling environment within the NMP niche. We show that Notch signalling influences early specification/differentiation of NMPs by steering them toward a presomitic/paraxial mesoderm fate at the expense of neurectoderm. *In vitro*, this appears to be mediated via a negative feedback loop between Notch and FGF signalling that is possibly critical for the proper calibration of the balanced production of neural and mesodermal cells from NMPs. Similar functional interactions between the two pathways have also been reported during the transition of axial progenitor-derived pre-neural and presomitic mesoderm cells toward spinal cord neurectoderm and somitic mesoderm respectively (Akai et al., 2005; Anderson et al., 2020; Diaz-Cuadros et al., 2020). Moreover, Notch signalling activity in the NSB/node embryonic regions at earlier stages of development was found to regulate progenitor cell contribution to the floor plate at the expense of notochord (Gray & Dale, 2010). Finally, we show that Notch signalling is also crucial for *HOX* gene activation in nascent NMPs during their induction from pluripotent cells, a cardinal hallmark of early posteriorisation of embryonic cells. This finding extends previous work linking control of *Hoxd* transcription and Notch signalling (Zákány et al., 2001). Our data suggest that Notch possibly exerts this role in NMPs through regulation of FGF signalling, a well-established driver of *HOX* gene transcription in the posterior growth region/axial progenitors (Delfino-Machín et al., 2005; Gogolou et al., 2022; Hackland et al., 2019; Mouilleau et al., 2021; van Rooijen et al., 2012). Moreover, Notch-mediated control of expression of some *HOX* genes appears to take place in a non-cell autonomous manner as indicated by their DAPT-sensitive transcriptional rescue in Notch-deficient/*TBXT* depleted hESC-derived NMPs upon co-culture with their wild-type counterparts. The role of the extrinsic environment in influencing cellular *Hox* codes has been pointed out previously with the demonstration that chick tail bud NMPs can switch from a Hox PG10+ to an “earlier” Hox PG8+ identity following transplantation into the NSB of younger host embryos (McGrew et al., 2008). We propose that Notch signalling is an integral part of the signalling environment within the NMP niche and a critical regulator of posterior body patterning.

## SUPPLEMENTARY FIGURES

**Figure S1.**
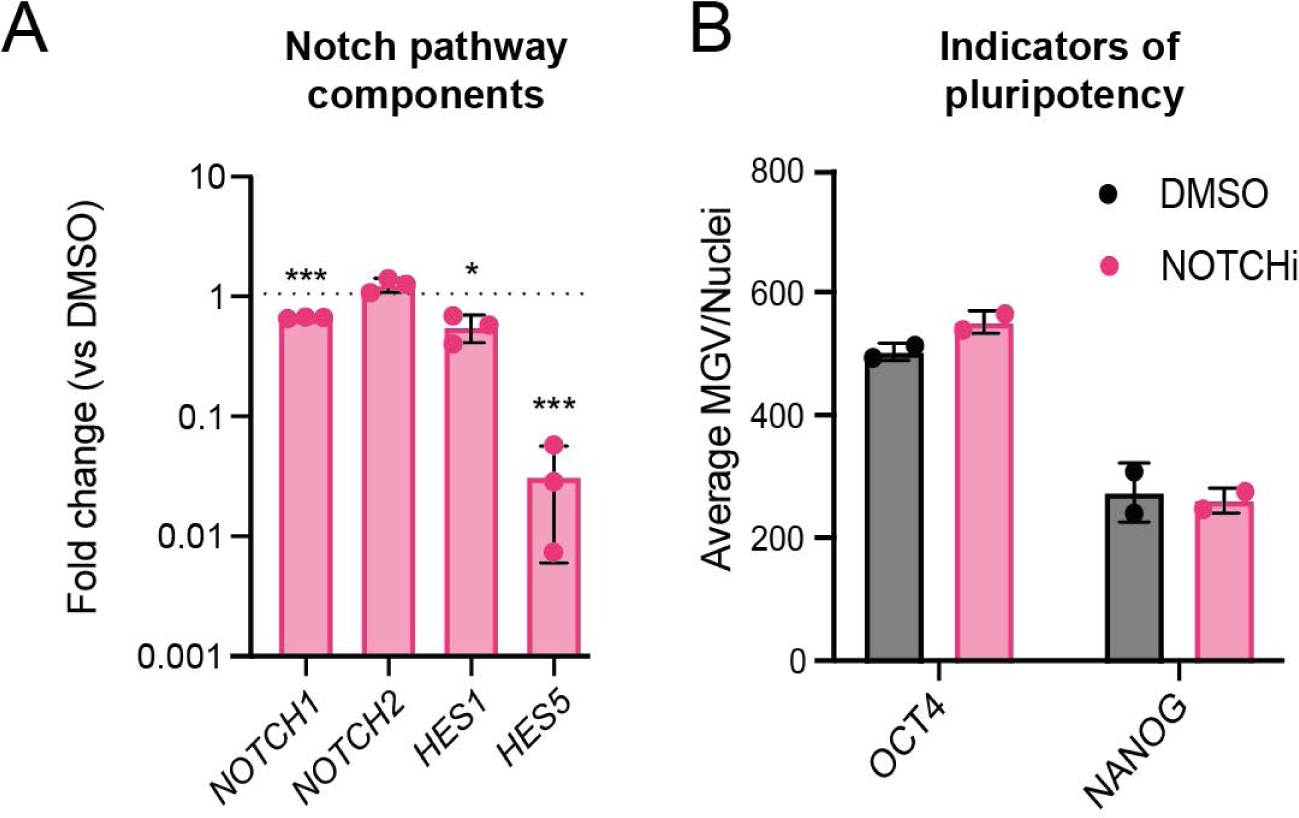
(A) qPCR expression analysis of indicated Notch signalling pathway components/targets in NOTCHi hESC-derived NMPs compared to DMSO controls. Error bars represent s.e.m n=3. *NOTCH*1 ***P<0.001, *HES1* *P=0.03, *HES5* ***P<0.001 (one sample t and Wilcoxon test). (B) Pluripotency associated marker expression in NOTCHi hESC-derived NMPs compared to DMSO controls. Error bars represent s.d. (n=2).

**Figure S2.**
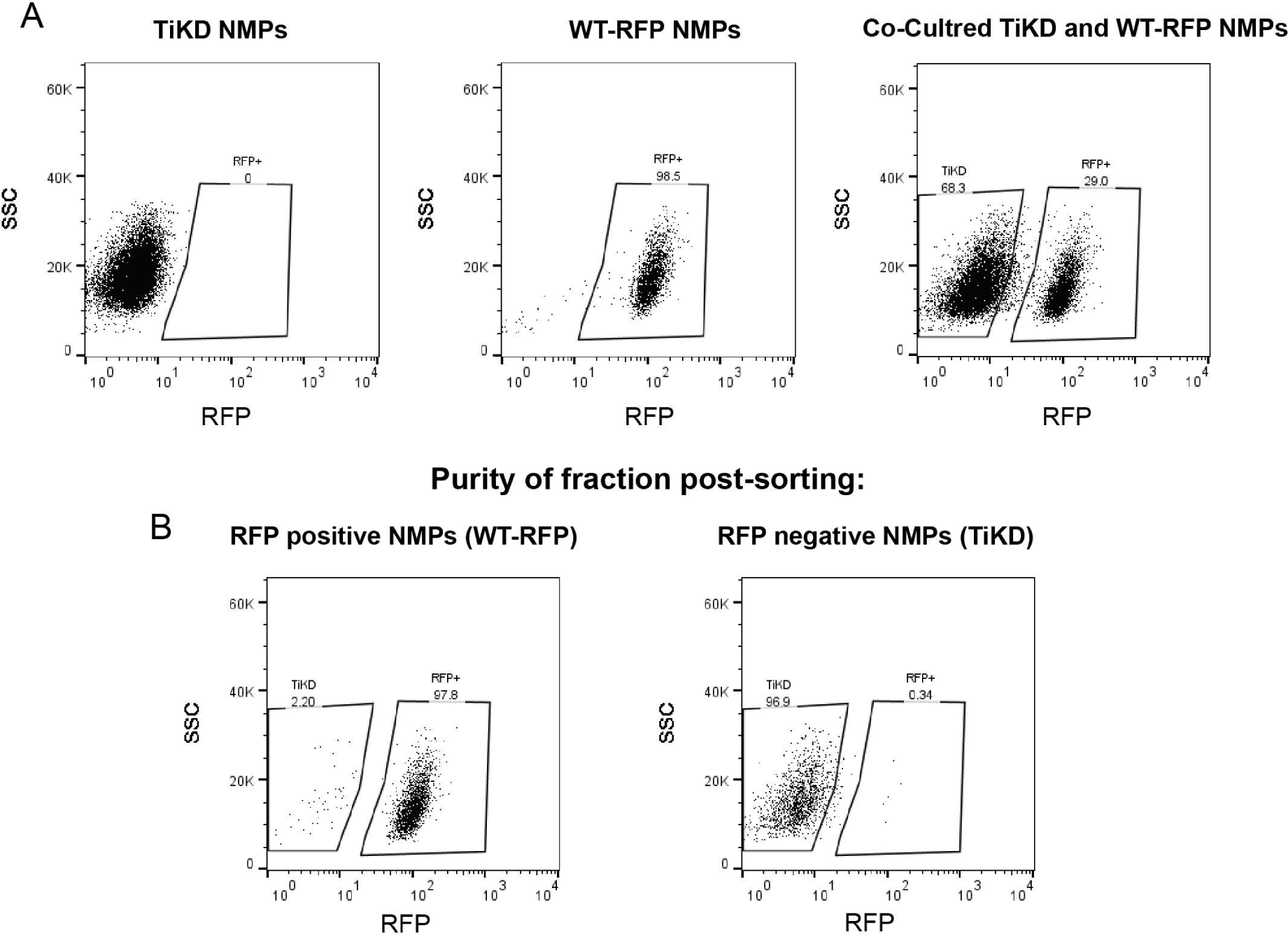
(A) FACS dot plots showing the fractions of RFP fluorescent reporter-positive cells in unlabelled TBXT knockdown (TiKD), wild type RFP (WT-RFP) hESC-derived NMPs and co-cultured (TiKD and WT-RFP) NMPs. (B) FACS dot plots showing the purity assessment following FACS of co-cultured NMPS into RFP negative (TiKD) and RFP positive (WT-RFP) fractions.

**Figure S3.**
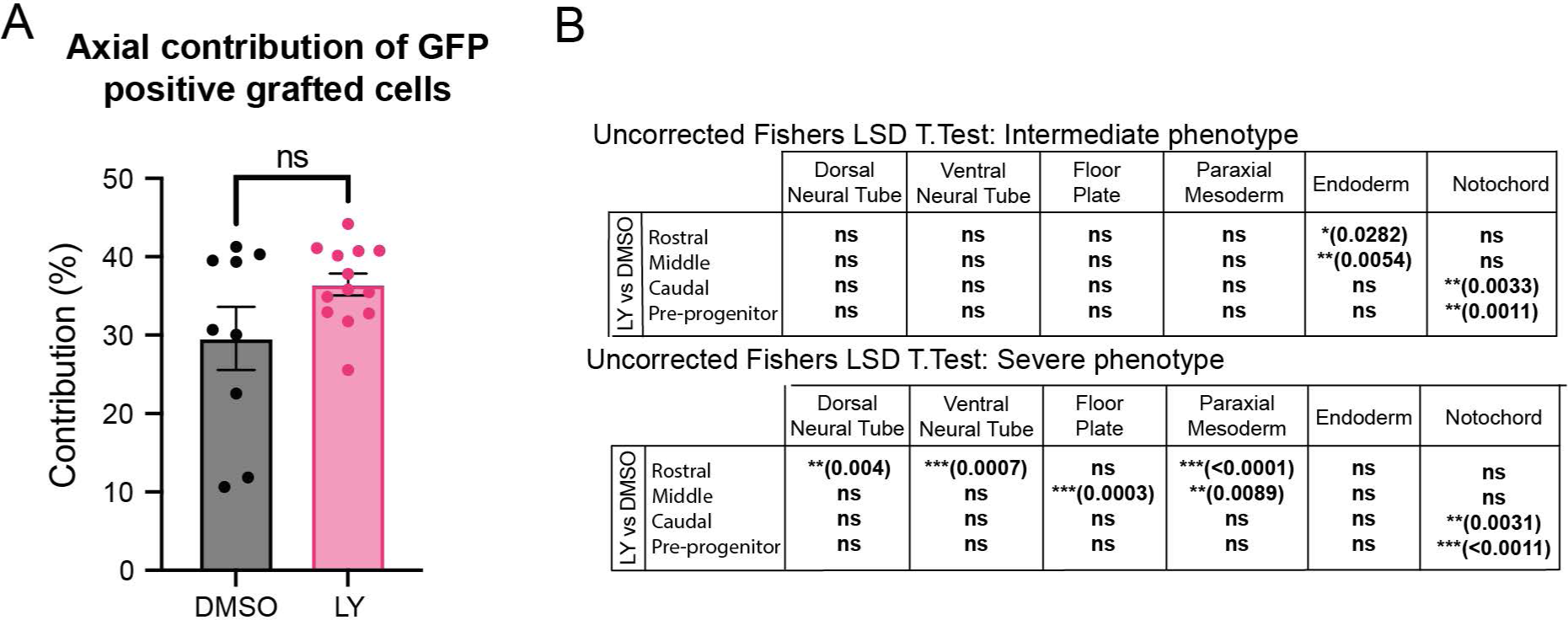
(A) Percentage anterior-posterior embryonic axis colonised by cells from the NSB following DMSO and LY treatment. Error bars indicate s.e.m (DMSO n=9 and LY n=13). ns P=0.07 (unpaired t.test). (B) Table showing the statistical P-value results for the severe and moderate LY phenotype using a one-way ANOVA (Fisher’s LSD test) (analysed sectioned embryos: DMSO n=9, LY severe n=4/9 and moderate n=5/9).

## MATERIALS AND METHODS

### Cell culture and differentiation

Use of hESCs has been approved by the Human Embryonic Stem Cell UK Steering Committee (SCSC15-23). The following hESC lines were employed: WA09 (H9), H9-RFP and *TBXT* shRNA sOPTiKD hESC lines (H9 background) (Bertero et al., 2016; Thomson et al., 1998). All cell lines were cultured routinely in feeder-free conditions in either Essential 8 (Thermo Fisher or made in-house) or mTeSR1 (Stem Cell Technologies) medium on Geltrex LDEV-Free reduced growth factor basement membrane matrix (Thermo Fisher). Cells were passaged twice a week after reaching approximately 80% confluency using PBS/EDTA or ReLeSR^TM^ (Stem Cell Technologies) as a dissociation reagent. *TBTX* inducible knockdown in the *TBXT* shRNA sOPTiKD hESC line was achieved using Tetracycline (Tet) hydrochloride (Merck Life Science) at 1 μg/ml as described previously (Bertero et al., 2016; Gogolou et al., 2022). hESCs were cultured in the presence/absence of Tet for 2 days prior to the initiation of differentiation and the Tet treatment was continued throughout the differentiation for the periods indicated in the results section/schemes. The RFP hESC line was generated following introduction of a pCAG-H2B-RFP plasmid (Price et al., 2021) into H9 hESCs using a 4D-Nucleofector (Lonza). After puromycin selection (1µg/ml), single cell deposition onto feeder cells was carried out followed by culture in 50% mTESR1:50% KnockOut™ Serum Replacement (Thermo Fisher) media, 20µM Cholesterol (Synthechol, Sigma), 10µM ROCK inhibitor. (Adooq Biosciences). The resulting clones were expanded, manually picked and cultured subsequently in mTeSR1. Cells were screened for mycoplasm using Lookout Mycoplasma PCR detection kit (Sigma-Aldrich) or Mycostrip detection kit (Invivogen). Cells were routinely screened for indicatiors of pluripotency OCT4, NANOG (Table S1) and SSEA4 (Adewumi et al., 2007; Draper et al., 2002).

For NMP differentiation, hESCs (70–80% confluent) were dissociated using Accutase solution (Merck Life Science) or TrypLE Select (Gibco) and plated at a density of 60,000 cells/cm^2^ on Vitronectin (Thermo Fisher) coated culture plates in N2B27 basal medium containing 50:50 Dulbecco′s Modified Eagle′s Medium (DMEM) F12 (Merck Life Science) / Neurobasal medium (Gibco) and 1 × N2 supplement (Gibco), 1 × B27 (Gibco), 1 × GlutaMAX (Gibco), 1 × Minimum Essential Medium Non-Essential Amino Acids (MEM NEAA) (Gibco), 2-Mercaptoethanol (50 μM, Gibco). The N2B27 medium was supplemented with CHIR (3 μM, Tocris), FGF2 (20 ng/ml, R&D Systems), and Rho-associated coil kinase (ROCK) inhibitor Y-27632 2HCl (10 μM, Adooq Biosciences) with the latter being withdrawn from the differentiation medium after the first day of NMP induction. DAPT (Tocris) was added at a concentration of 50 µM and DMSO was used at 5 µl/ml as control. PD032590 (Merck) was used at 1 μM. For *TBXT* inducible knockdown, NMP medium was supplemented with 1 μg/ml Tet hydrochloride and replenished every other day.

### Flow cytometry

After co-culture of 50% unlabelled TiKD and 50% RFP+ wild type hESCs and differentiation towards NMP, unlabelled NMPs were sorted at day 3 of differentiation using a FACS Jazz cell sorter (BD). Gates were set using unlabelled and RFP+ cells independently. Purity checks were done post sort. Data were analysed with FlowJo software (BD) (See Figure S2).

### Immunofluorescence and imaging

Cells were fixed in 4% Paraformaldehyde (PFA) for 10 min at room temperature, rinsed twice with PBS and permeabilised/blocked with blocking buffer containing 0.1% Triton X-100 in PBS containing 1% bovine serum albumin (BSA) for 1-2hr at room temperature (RT). Primary antibodies were diluted in the blocking buffer and cells were incubated with primary antibodies overnight at 4°C. Following three washes with PBS, cells were incubated with secondary antibodies conjugated to Alexa fluorophores (Invitrogen) diluted in blocking buffer for 2-4 hr at RT, in the dark. Cell nuclei were counterstained with DAPI:PBS (Thermo Fisher, 1:12000) and fluorescent images were acquired using the InCell Analyser 2200 system (GE Healthcare). Images then were processed in Fiji (Schindelin et al., 2012) using identical brightness/contrast settings to allow comparison between different treatments. The positive/negative threshold (75^th^ percentile) was set using a sample incubated with secondary antibody only. Antibodies and corresponding dilutions are shown in Table S1.

### Western blotting

Pelleted cells lysed in RIPA lysis buffer (50 mM Tris-HCl pH8.0, 100 mM NaCl, 2 mM MgCl2, 1 % Triton X-100, 0.1 % sodium deoxycholate, 0.1 % SDS supplemented with 1 mM DTT, 1× Complete protease inhibitor cocktail (Roche) and 250 U Benzonase nuclease immediately before use) for 10 mins at 37°C followed by centrifugation to remove insoluble debris. 50 µg of protein lysate per lane was then run on a NuPage 4-12% Bis-Tris gel (Thermo Fisher) at 120 V. Proteins were then transferred to a nitrocellulose membrane (Trans-Blot Turbo Mini 0.2 µm Nitrocellulose Transfer) using Trans-Blot Turbo Transfer System (Bio-Rad) following manufacturers guidelines. Membranes were then wash in TBS-T and blocked in 5% BSA: TBS-T for 1hr at RT. Membrane was incubated with primary antibodies (Table S1) overnight at 4°C followed by HRP-conjugated secondary antibodies for 1hr at RT. ECL detection was enhanced using SuperSignal West Pico PLUS (Thermo Fisher) as per the manufacturers guidelines and imaged using a G:BOX Chemi XX98 imager (Syngene). Images then were processed in Fiji (Schindelin et al., 2012).

### Quantitative real time PCR

Total RNA was extracted using the total RNA purification kit (Norgen Biotek) following the manufacturer’s instructions. The cDNA sysnthesis was completed using the High-Capacity cDNA Reverse Transcription kit (Thermo Fisher). Quantitative real-time PCR was carried out using the QuantStudio 12 K Flex (Applied Biosystems) thermocycler in combination with the Roche UPL system and the TaqMan Fast Universal PCR Master Mix (Applied Biosystems) or with PowerUp SYBR master mix (Thermo Fisher). Primer sequences and corresponding probes (where applicable) are shown in Supplementary Table S2. Graphs were generated using GraphPad Prism (GraphPad Software), which was also employed for statistical analysis.

### Chick embryo grafting experiments

White Leghorn *Gallus gallus* (eggs obtained from Henry Stewart & Co., Lincolnshire and Winter Farm, Royston) or GFP-expressing chick embryos [Roslin Institute, Midlothian (McGrew et al., 2004) were incubated until Hamburger Hamilton (HH) stage 4 and then dissected from the egg and cultured in vitro until HH8. Embryos were cultured on media plates containing either a γ-secretase inhibitor dissolved in the solvent dimethyl sulfoxide (DMSO) or on media plates containing DMSO alone. The concentration of LY411575 γ-secretase inhibitor (made in-house, University of Dundee) used was 150nM. Embryos were transferred to fresh culture plates every 12 hours to maintain optimal inhibitor activity. Following in vitro culturing the NSB region from HH8 GFP transgenic donor chicks was isolated and grafted to a homotopic location on stage matched wild-type donor embryos. Embryos were then returned to in vitro culture plates for a further 27 to 29 hours to allow for progenitor cells within the NSB to contribute to axial and paraxial tissues. Subsequently, embryos were fixed, cryosectioned and analysed by cell count for tissues that were colonised by GFP-positive cells across the rostral, middle, caudal and pre-progenitor domains. Each embryo had 5 sections from each axial domain analysed by cell count analysis in each domain. The proportion of counted cells in a particular tissue from one section was scored as a proportion of the total GFP-positive cells in that section. The proportion of cells in a particular section was used for analysis as opposed to the raw values obtained so as to exclude variation in cell number between sections and embryos from biasing the analysis. The proportion data on GFP-positive cells in axial and paraxial tissues were pooled between embryos of the same treatment group and axial domain to obtain a mean value. These values therefore represented the mean proportion of cell contribution to specific tissues at specific anterior-posterior axial locations. Pairwise comparisons were made between the GFP cell counts of LY and DMSO treated embryos in each cell type at each of the rostral, middle, caudal and pre-progenitor domains and were subjected to statistical tests to determine where significant differences occurred.

## ACKNOWLEDGEMENTS

We would like to thank Prof. Ivana Barbaric (University of Sheffield) for providing the H2B-RFP expression vector. We are grateful to Matt French, Sally Lowell, Matt Towers and Val Wilson for critical reading of the manuscript.

## COMPETING INTERESTS

The authors declare no competing or financial interests.

## AUTHOR CONTRIBUTIONS

Conceptualization: AT, FC, JKD; Formal analysis: FC, CS, SH, SG; Investigation: FC, CS, SH, AG, SG, TF, DS, KB, BMJ; Resources: AG, TF, DS; Writing – original draft preparation: FC, AT; Writing – review and editing: FC, CS, SH, AG, TF, DS, BMJ, KB, DB, JKD, AT; Visualization: FC, CS, AT, SH, JKD; Supervision: AT; Project administration: AT; Funding acquisition: JKD, DB, AT.

## FUNDING

This work was supported by funding from the Biotechnology and Biological Sciences Research Council (BB/P000444/1), the European Union Horizon 2020 Framework Programme (H2020-EU.1.2.2; grant agreement ID 824070) and the Medical Research Council (MR/V002163/1) to AT. KB was supported by a White Rose BBSRC Doctoral Training Partnership (DTP) in Mechanistic Biology studentship (BB/T007222/1). SG was supported by an MRC New Investigator award to JKD (G0400349: Analysis of primitive streak stem cells and the role of Notch in their axial mesoderm derivatives).

## SUMMARY STATEMENT

Notch signalling is a critical regulator of the induction and differentiation of posteriorly-located neuromesodermal axial progenitors, the precursors of the neural and mesodermal components of the amniote embryonic body trunk.

## Notes

### Competing Interest Statement

The authors have declared no competing interest.

